# Nanoparticle-induced biomembrane fusion: unraveling the effect of core size on stalk formation

**DOI:** 10.1101/2023.05.22.541080

**Authors:** Giorgia Brosio, Giulia Rossi, Davide Bochicchio

## Abstract

Membrane fusion *in vitro* is a strategy to load model or cell-derived vesicles with proteins, drugs, and genetic materials for theranostic applications. It is thus crucial to develop strategies to control the fusion process, also through synthetic fusogenic agents. Ligand-protected, membrane-penetrating gold nanoparticles (Au NPs) can facilitate membrane fusion, but the molecular mechanisms remain unresolved. Here, we tackle NP-induced stalk formation using a coarse-grained Molecular Dynamics approach and enhanced sampling techniques. We show that smaller (2 nm in diameter) NPs lead to a lower free energy barrier and higher stalk stability than larger NPs (4 nm). We demonstrate that this difference is due to a different ligand conformational freedom, which in turn depends on the Au core curvature. Our study provides precious insights into the mechanisms underlying NP-mediated membrane fusion, while our computational approach is general and applicable to studying stalk formation caused by other fusogenic agents.

**TOC GRAPHICS:** 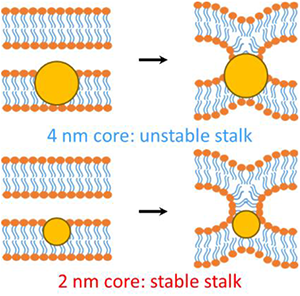

---

Membrane fusion is fundamental for many biological activities such as membrane trafficking^1^, synaptic transmission^23^, fertilization^4,5^, and viral infection^6^. It is a complex process involving the concerted movement and rearrangement of many different molecules (mainly lipids and proteins). The most likely fusion path, from two approaching bilayers to the opening of the fusion pore, includes intermediate metastable minima (Figure 1a). At first, the two separated membranes must be brought close to each other, typically at < 5 nm distance. Then, a point-like protrusion of a lipid tail can initiate the hydrophobic contact between the membranes, leading to the so-called stalk, a configuration allowing the mixing of the lipids from the outer leaflets^7–10^. Stalk elongation may lead to a second metastable configuration, the hemifusion diaphragm, and then the process is completed with the opening and expansion of a fusion pore^10–12^. The fusion pore opening could directly evolve from the stalk configuration.^12^ The different minima characterizing this complex multi-step process can be separated by relatively high free energy barriers^8,9,13^ (Figure 1b).

**Figure 1:**
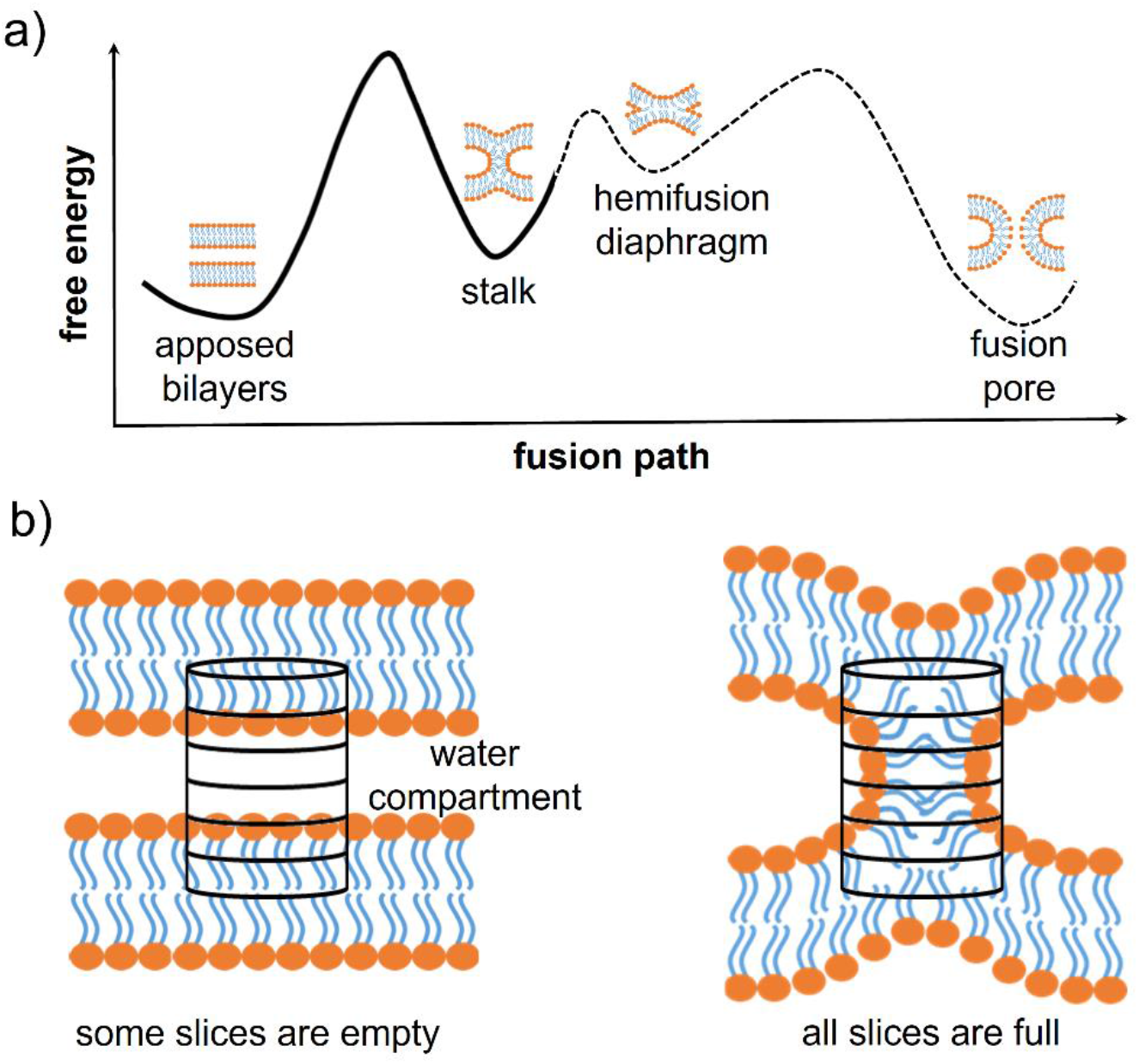
fusion intermediates and definition of the chain coordinate. a) Scheme of the typical free energy landscape along the fusion path. It exhibits three free energy barriers: one for the stalk formation, one for the expansion of the hemifusion diaphragm, and one for the opening of the fusion pore. b) A scheme of the cylinder used by Hub and co-workers to define the chain coordinate ξ_ch_.^9^ The cylinder is divided into slices and placed in between two adjacent membranes. When the membranes are distinct, most slices are empty of lipid tails, while in the stalk configuration, all slices are filled

*In vivo*, overcoming the fusion barriers is generally achieved thanks to the presence of specialized proteins, such as the SNARE complex, which catalyze the fusion of synaptic vesicles with the plasma membrane^12,14^. *In vitro*, achieving controlled fusion by means of artificial fusogens would be critical to realize artificial vesicles with controlled membrane composition and cargo^15,16^ Gold nanoparticles (Au NPs), functionalized with anionic (11-mercapto-1-undecanesulfonate – MUS) and neutral, hydrophobic (octanethiol – OT) ligands, are promising synthetic fusogenic agents. Indeed, a recent study by Tahir *et al*. demonstrated that MUS:OT NPs induce fusion between pure phosphatidylcholines(PC) liposomes^17^, and our group showed that the liposome cholesterol content can finely tune MUS:OT NP-induced fusion events. ^18^

Molecular Dynamics (MD) simulations, thanks to their high spatiotemporal resolution, have been successfully employed to shed light on the molecular mechanisms involved in the different stages of the fusion process, either spontaneous ^19–22^ or mediated by fusion agents^17,23–25^. The presence of high free energy barriers makes the use of atomistic models challenging and calls for coarse-grained models and/or enhanced sampling techniques. Thanks to coarse-grained modelling, spontaneous NP-induced stalk formation has been observed in unbiased MD, but only increasing the temperature much above the physiological one. ^18^ Therefore, the derivation of the free energy profile of stalk formation requires enhanced sampling, and the challenge is the definition of a collective variable (CV) that approximates well the reaction coordinate. Finding a CV suitable to describe stalk formation is arduous because of the high number of molecules involved and their high conformational freedom, and a poor CV could easily lead to hysteresis in the free energy calculation along the stalk-formation and stalk-destruction pathways. ^26^

The group of J. S. Hub recently solved these issues by proposing the so-called “chain coordinate” ξ_ch_, ^9,26–28^ designed to study stalk formation between two flat lipid bilayers and implemented directly into the Gromacs MD suite. In order to quantify the degree of connectivity between the hydrophobic cores of two lipid membranes separated by a water layer, one can imagine to track the filling/unfilling of a cylindrical volume placed in between the two membranes, with its axis aligned to the membrane normal, as shown in Figure 1c. ξ_ch_ represents the cylinder’s degree of filling by hydrophobic lipid tails. In the stalk configuration, all slices are filled with hydrophobic lipid tails, and ξ_ch_ approaches unity, while when the membranes are separated, several slices are empty, and ξ_ch_ is << 1 (see SI for further details on the ξ_ch_ original definition). Hub and co-workers exploited ξ_ch_ in umbrella sampling calculations and showed how lipid membrane composition impacts the stalk formation free energy path^9^.

Here we build on the original definition of the ξ_ch_ chain coordinate, modifying it to account for the presence of the fusogenic agent, a MUS:OT Au NP embedded in one of the two lipid bilayers. Our previous MD simulations based on a coarse-grained force field revealed that the stalk rim constantly forms over a single Au NP^18^, consequently to the formation of a hydrophobic contact between a lipid tail of the top bilayer and transient hydrophobic defects on the NP, water-facing surface. This mechanism is consistent with the one described in the literature for NP-free membranes, with the only difference being that the contact is a lipid-NP one and not a lipid-lipid one^9^. Therefore, we decided to redefine the chain coordinate ξ_ch_ to make it suitable to deal with a NP-induced stalk formation (Figure 2), implementing the following two changes.

**Figure 2:**
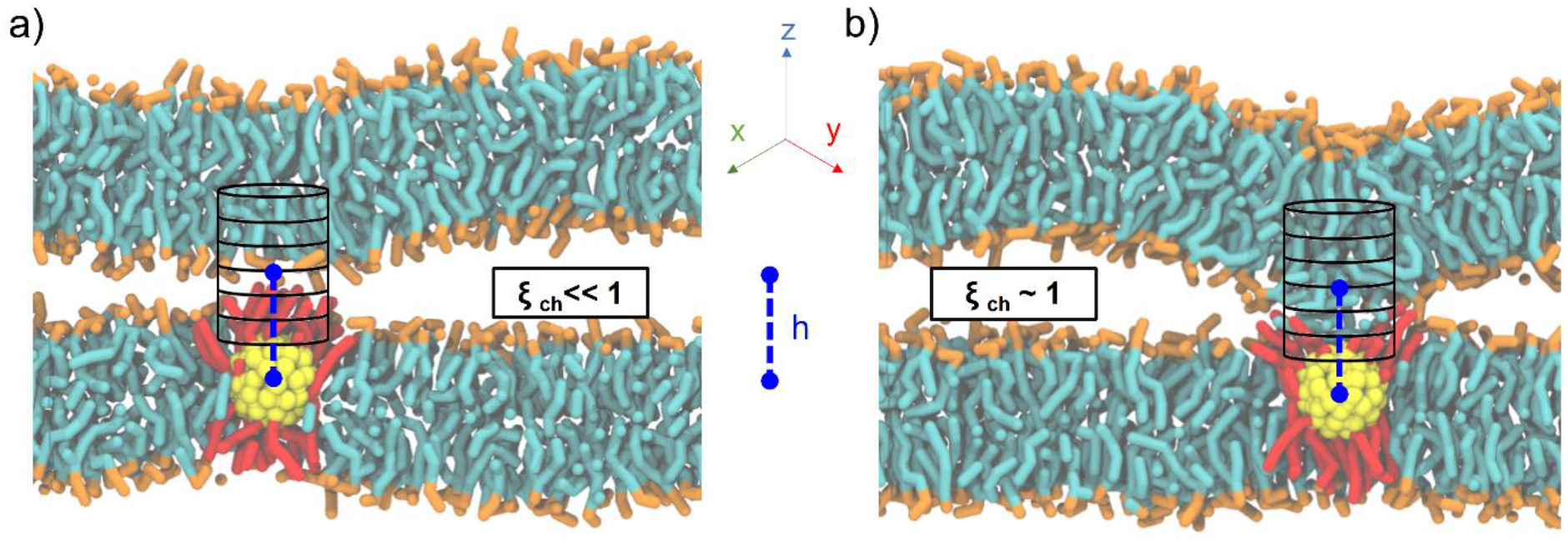
a new implementation of the chain coordinate. The cylinder is now defined with respect to the NP, following it in the *xy* plane and being located along *z* at a distance *h* from the NP center of mass. a) When the membranes are still distinct, the cylinder is partially empty (ξ_ch_<< 1). b) When a stalk is formed over the NP, the cylinder is almost full of hydrophobic beads (ξ_ch_ ∼ 1). Lipid heads are in orange, lipid tails in cyan, the NP core in yellow, MUS ligands in red, OT ligands and cholesterol are not shown for clarity.

First, to have the cylinder always follow the NP diffusing in the membrane plane, we set the in-plane *x* and *y* coordinates of the cylinder axis to coincide with the *x* and *y* of the NP center of mass (COM). Second, to account for the system asymmetry along the membrane normal and the possible differences in NP size, we set the *z* of the cylinder center of mass to a user-defined distance *h* from the NP COM *z* coordinate (see SI for more details). As shown in Figure 2, even in the presence of NP diffusion and oscillation along the *z* direction, the cylinder is automatically centered over the NP, thus monitoring the degree of hydrophobic connection established between the two lipid bilayers just on top of the NP.

To test our newly defined CV, we set up a system consisting of two DOPC bilayers with 30% mol cholesterol, a lateral size of ∼19 nm, and a single NP with a 2 nm core fully embedded in the lower bilayer (see Supporting Figure S1a). After minimization of our solvated system, we equilibrated it and then derived stalk formation free energy profiles along ξ_ch_ using the Umbrella Sampling (US) technique. We extracted the starting configuration of each of the 16 windows from a simulation in which we slowly pulled the system along ξ_ch_ to drive stalk formation (see SI for more details). We set the shift parameter *h* to 2.75 nm to account for the size of the NP core. The free energy profile (reported in Figure 3a) was constructed from the umbrella histograms with the weighted histogram analysis method (WHAM), ^29^ as implemented in the *gmx wham* Gromacs tool, and the error was calculated from a bootstrap procedure. ^30^

**Figure 3:**
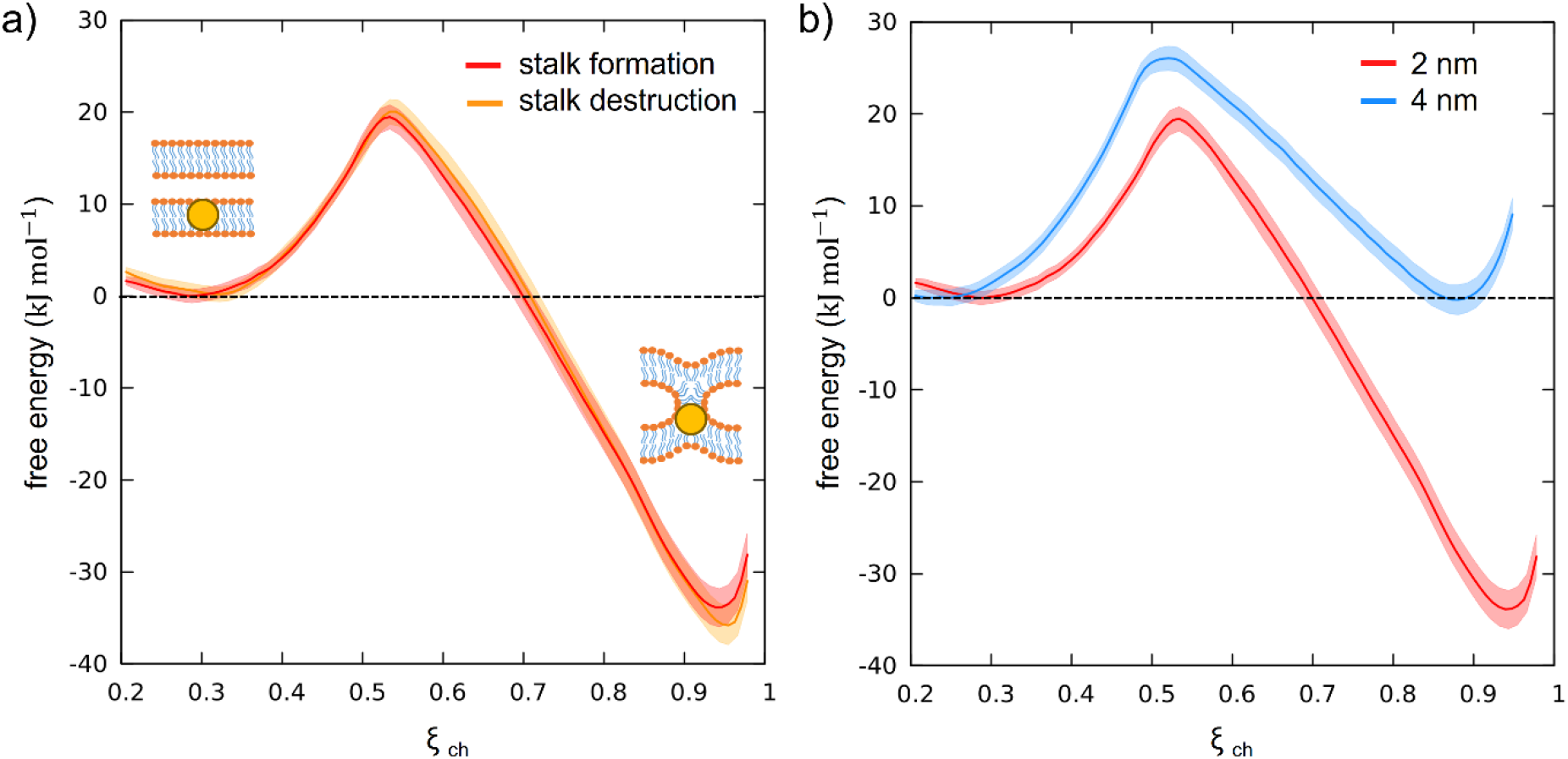
free energy profiles of stalk formation. a) Free energy profiles of stalk formation (red) and destruction (orange) as a function of ξ_ch_ for NP with a core size of 2 nm. The absence of hysteresis demonstrates the reliability of the CV. b) Free energy profiles of stalk formation for a 2 nm core (red) and a 4 nm core (blue). The barrier to stalk formation in the second case is higher, while the stalk becomes much less stable, resulting in a minimum with the same free energy as the apposed membranes one.

The free energy profile, shown in red in Figure 3a, shows a local minimum at ξ_ch_=0.31 that corresponds to the state in which the two membranes are separated (pre-stalk state). The second minimum, at ξ_ch_ =0.94, corresponds to the state in which the stalk is fully formed. In this case, the stalk is thermodynamically favored, being lower in free energy of about 30 kJ mol^-1^, and the barrier to stalk formation is about 20 kJ mol^-1^. This picture is consistent with previous unbiased simulations, which were run at a higher temperature and estimated a barrier of about 15 kJ mol^-1^ in the presence of cholesterol.^18^ Our calculation here demonstrates that, at physiological temperature, the stalk formed on top of the NP is highly stable, with a barrier to stalk destruction larger than 50 kJ mol^-1^.

To test the consistency of our method and exclude the presence of significant hysteresis, we calculated the same free energy profile using initial configurations taken from a reverse pulling simulation, in which the system has been brought from the stalk state back to the pre-stalk state. We refer to this second profile as the “profile of stalk destruction” (Figure 3a, orange). In the ideal case, if the free energy profiles are at convergence, the free energy profiles of stalk formation and destruction should be identical. Comparing the two profiles in Figure 3a, it can be noticed that they overlap within the error bars, proving the absence of hysteresis and thus the reliability of our CV. Once verified the appropriateness of our method, we tackled the effect of the NP core size on the stalk formation process. To this purpose, we constructed a second system, still containing two membranes composed of 70% DOPC and 30% cholesterol, but with a single, embedded NP with a gold core diameter of 4 nm (see Supporting Figure S1 b.). The free energy profile of stalk formation was calculated as in the previous case (see SI for more details), with the only different parameter being the *h* shift, which was increased to 3.75 nm to account for the new core radius. Once again, we verified the absence of significant hysteresis by calculating the free energy profile of stalk destruction, and obtained an almost perfect overlap with the one of stalk formation (see Supporting Figure S2).

Figure 3b compares the free energy profiles of stalk formation in the two systems. We can notice a different barrier and a different depth of the stalk minimum. In the presence of the bigger NP, the stalk is not thermodynamically favored anymore, becoming equally probable to the pre-stalk state. Furthermore, the barrier to stalk formation is higher, suggesting a slower kinetics of stalk formation for the large NPs. This result is consistent with recent experimental observations in which MUS:OT NPs with a core size of ∼2.5 nm promoted hemifusion between adjacent bilayers, while larger NPs exhibited only surface binding to membranes. ^31^

In order to analyze and explain the molecular origin of this size effect, we calculated and compared a few crucial quantities in both systems, relating them to the differences observed in the free energy profiles of Figure 3b.

The molecular trigger for stalk formation is the establishing of a first contact between the hydrophobic tail of a lipid protruding from the facing bilayer (in our set-up, the one above the NP) and a transient hydrophobic defect in the NP ligand shell that is exposed to the water phase (see Supporting Figure S3). This is a rare event, whose rate depends on the height of the free energy barrier separating the pre-stalk state from the stalk state. We wanted to test the hypothesis that the more hydrophobic surface is exposed by the NP to the aqueous environment, the more likely it is that a hydrophobic NP-lipid contact is established. To quantify this, we defined the H-LASA as the hydrophobic lipid-accessible-surface-area of the NP divided by the total lipid accessible surface area of the NP (see SI for more details). We then monitored the time evolution of the H-LASA during a simulation (see Supporting Figure S4), and recorded its maximum value.

Figure 4a shows the maximum H-LASA obtained for NPs of both sizes when embedded in the membrane in the pre-stalk configuration. The maximum H-LASA anti-correlates with the barrier to stalk formation shown in Figure 3, being significantly larger for the 2 nm core size NP than for the 4 nm one. This difference can be explained with simple geometric considerations. As the core radius becomes bigger, the average surface curvature decreases, while the ligand grafting density and ligand length remain the same. Therefore, the interface of the large NP will have both a higher ligand density and a higher surface charge density than a smaller NP. Both the ligand density and the charge density limit the ligand conformational freedom and make the formation of hydrophobic patches more difficult. The same effect is present when the NPs are solvated in water (see Supporting Figure S5), and it is responsible for hindering the hydrophobic aggregation between the large NPs. ^32^

**Figure 4:**
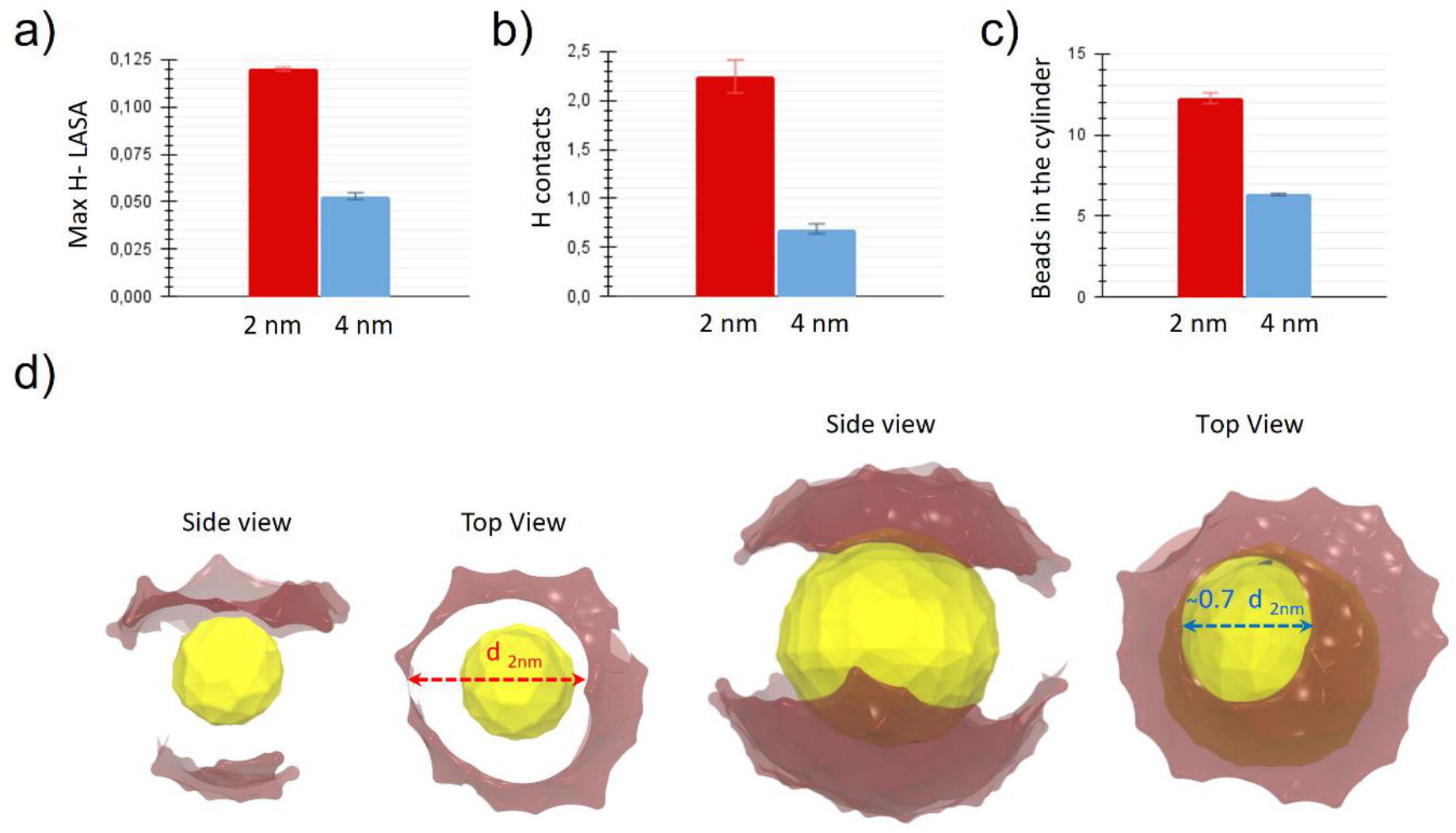
microscopic interpretation of the NP core size effect on stalk formation. a) Maximum of H-LASA in the pre-stalk configuration for the NPs with a diameter of 2 nm and 4 nm. b) Number of hydrophobic contacts in the stalk state for the two NPs. c) Number of hydrophobic lipid beads present in the cylinder in the stalk state for the two different size NPs. d) Representative snapshots with views from the side and from the top of the NPs of the two sizes in the stalk state. The opening of the MUS-charged terminals due to the formation of the stalk above the nanoparticle is highlighted. We show only the MUS charged terminals (in transparent red) and the NPs’ core (in yellow). The width of the “hole” corresponds to the diameter of the stalk above the NP, which is different in the two cases.

The impact of the NP core size on the stalk stability is even more pronounced than the effect on the barrier to stalk formation. What is the molecular origin of the higher stalk stability in the case of the small NP? A stalk is a hydrophobic connection between two membranes, and we expect its stability to be correlated with the stalk geometry and composition. A larger/denser stalk is expected to provide a more stable hydrophobic connection between the membranes, even when mediated by the NP. Are the stalk dimensions and densities different in the two systems investigated here?

We calculated the number of hydrophobic contacts between the NP and the lipids in the stalk region. More precisely, we calculated the variation of the number of contacts between the hydrophobic beads of the NP ligands and those of the membranes between the pre-stalk and the stalk state. The results, shown in Figure 4b, correlate well with the different stalk stability. In the case of the 2 nm NP, the system takes advantage of the possibility of forming more NP-lipid hydrophobic contacts and thus manages to create a more stable stalk.

To get a quantitative estimation of the stalk diameters, we counted all the hydrophobic beads contained in the 18 slices of the cylinder closer to the NP and thus located in the central part of the stalk region (see SI for more details) . The results are shown in Figure 4c. The stalk contains more hydrophobic beads when the NP is smaller. From the values obtained, being the cylinder height constant, it is straightforward to derive the ratio between the average diameter of the stalk in the case of a 4 nm core and the one in the case of a 2 nm core, which turns out to be about 0.7. The difference in stalk diameter between the two systems becomes evident in Figure 4d, where we show representative snapshots of the NPs in the stalk state. The opening of the charged MUS terminals due to the formations of the stalk above the NP is highlighted by our representation, where the diameter of the “hole” corresponds to the diameter of the hydrophobic connection between the NP and the membrane above. The hole (and thus the stalk) is larger in the system containing the small NP. Again, the molecular origin of this difference is the higher conformational freedom of MUS ligands on the small NP.

In conclusion, we developed a reliable procedure to calculate the free energy of stalk formation induced by functionalized NPs. Our approach was based on adapting the chain coordinate^9^ to the presence of membrane-embedded fusogenic agents. We calculated the free energy profiles of stalk formation via umbrella sampling, revealing that the NP core size can significantly impact both the kinetics and the thermodynamics of the process. A smaller NP core leads to higher conformational freedom of charged ligands, which can more easily rearrange to allow for the first hydrophobic contact between the NP and the facing lipids, a contact that triggers stalk formation. Coherently, when the core is smaller, the open-ligand configuration is more stable, and it allows for the formation of a larger, durable stalk, driving its thermodynamic stability over the pre-stalk state.

Stalk formation is a fundamental step of membrane fusion, and thus our results have important implications for the design of artificial fusogenic agents. In particular, we highlight that ligand flexibility and conformational freedom are critical features that can be exploited to increase the efficiency of the fusion process. The biased protocol developed here will be used to study the influence of the membrane composition on NP-induced stalk formation and will be combined with further *in silico* studies to reveal how the system can go beyond the stalk state, completing the fusion process. It is worth noting that our collective variable definition is quite general and, thus, not limited to the particular type of NP studied herein. Indeed, our protocol could be easily adapted to stalk formation induced by different artificial or biological agents, helping develop new strategies to induce and control fusion events.

## ASSOCIATED CONTENT

The following files are available free of charge:

Supporting Information.pdf

## AUTHOR INFORMATION

The authors declare no competing financial interests.

## ACKNOWLEDGMENT

The authors acknowledge funding by MIUR – DIFI Dipartimento di Eccellenza 2018-2022 for computational resources.

## Notes

### Competing Interest Statement

The authors have declared no competing interest.

