## Supporting Information for "Nanoparticle-induced biomembrane fusion: unraveling the effect of core size on stalk formation"

### Electronic Supplementary Information

AUTHOR INFORMATION

**Corresponding Author**

*Davide Bochicchio,.

**Simulation set-up**

To study the effect that the core size of a nanoparticle has on the formation and stability of the stalk state we used the two double membrane systems shown in Figure S1. In Figure S1a and S2b a MUS:OT AuNP with two different core size (2nm and 4nm) are fully embedded in the lower bilayer, inserted in such a way that their ligands are in contact with the upper membrane.

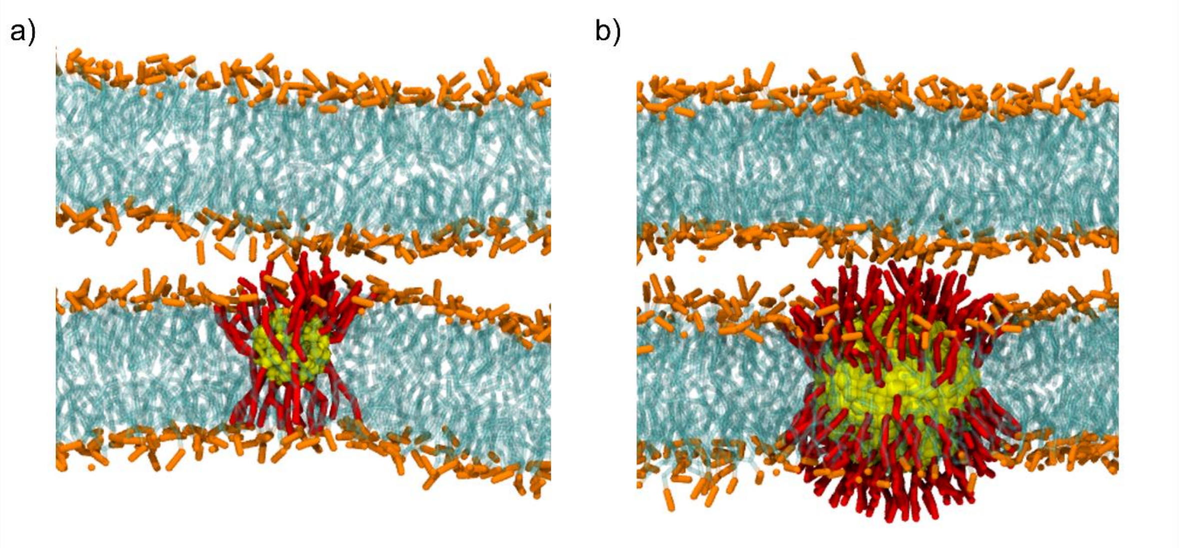


**Figure S1: the two double membrane and NP systems**. Snapshot representing two systems, composed of two parallel DOPC + 30% mol cholesterol. a) A single 70:30 MUS:OT NP with core radius of 2nm is embedded into the bottom bilayer. b) A single 70:30 MUS:OT NP with core radius of 4nm is embedded into the bottom bilayer. DOPC is in cyan, nanoparticle core in yellow and the lipid heads in orange. MUS ligands are shown, in red. The solvent and OT ligands are not shown for clarity.

We started with a box containing a single membrane and a fully embedded NP: the NP lies at the center of the bilayer and the charged ligands are distributed between the entrance and the distal leaflet. A second box of the same size was created, containing another membrane NP-free, with the same lipid composition. Both systems were initialized with the INSANE software. [1] To allow for a meaningful comparison, in both systems the lateral size of the simulated membranes has been in the range of 19 nm, and the amount of water in between the two membranes has been set to obtain a comparable hydration (around 7 water beads per DOPC head).

Before being merged in a single box, the two membranes (one with the NP and one without) were minimized and equilibrated separately. Then the minimization and equilibration were repeated. The solvent used for the simulations is water, but for neutralizing the system and mimicking the physiological salt concentration we added sodium, chloride and calcium beads.

All the previous simulations were carried with GROMACS software package, [2] in the NpT ensemble and with periodic boundary conditions (PBC) in all directions. The temperature was set to T = 310 K using the v-rescale thermostat [3] (with a time constant t = 1 ps). The pressure was controlled by the Parrinello-Rahman barostat[4] in the production runs, while we used the Berendsen barostat [5] in the equilibration runs and in the production runs for which Parrinello-Rahman was unstable. A semi-isotropic pressure coupling was used (the *xy* directions were coupled independently of the *z* direction), allowing for spontaneous membrane deformations.

Interaction potentials were described with the coarse-grained Martini force field[6]. As regards the *in silico* model of the 30:70 MUS:OT NPs used in this work, it had been developed and validated in previous works. [7–10]

**Reaction coordinate for stalk formation**

Our starting point was the CV recently proposed by the group J. S. Hub, named "chain coordinate", $\xi_{ch}$, designed to study the stalk formation in the absence of external agents, neither synthetic nor protein, and implemented directly into the pull code of Gromacs 2018.8.

The chain coordinate is defined thanks to a cylinder divided into $N_s$ slices located in the aqueous medium between two membranes. In particular, $\xi_{ch}$ is defined as the fraction of slices filled by hydrophobic beads belonging to the lipid tails. The cylinder has radius $R_{cyl}$, and its slices (each with thickness $d_s$) are symmetrical with respect to the center of mass $z_{mem}$ of the hydrophobic membrane atoms along the membrane normal. In this way, the center of the slice s is $z_s = z_{mem} + (s + 1/2 - N_s/2)d_s$ with s ranging from $0$ to $N_s - 1$. $\xi_{ch}$ can thus be expressed as:

$$\xi_{ch} = N_s^{-1} \sum_{s=0}^{N_s-1} \delta_s \, n_s^{(ap)}$$

where $n_s^{(ap)}$ is the number of hydrophobic atoms within slice $s$ of the cylinder. Both hydrophobic lipid tail beads and hydrophobic beads of cholesterol were used as beads contributing to $\xi_{ch}$. $\delta_s$ is a continuous indicator function ($0 \leq \delta_s < 1$), which equals zero if no hydrophobic atom is in slice s ($n_s^{(ap)} = 0$) and tends to unity if one or multiple hydrophobic atoms are located in the slice ( $n_s^{(ap)} \geq 1$).

From previous unbiased simulations, [11] carried out using a temperature of 370 K, it was observed that in the presence of a NP the stalk rim always forms on top of it, consequently to the formation of a hydrophobic contact between a lipid tail of the facing bilayer and hydrophobic beads of the NP ligands (see Figure S3a).

Our first step to calculate a reliable free energy profile has been the redefinition of Hub chain coordinate $\xi_{ch}$to take into account the presence of a single MUS:OT nanoparticle. To this purpose, we had to edit the Gromacs source code so that the center of the cylinder was located at a user-defined distance *h* along the *z*-axis from the NP center of mass (COM). We redefined the center of the slice *s* along *z* as is $z_s = z_{NP} + (s + 1/2 - N_s/2)d_s + h$ where $z_{NP}$ is the center of mass of NP along the membrane normal. In this way, the cylinder is always placed above the NP and thus can be used to force the presence or the absence of a stalk in that region, which is where the stalk form in unbiased simulations. Furthermore, if the NP diffuses in the *xy* plane, the cylinder automatically follows it, allowing the stalk to remain always over the NP.

**Implementation of the code**

We modified a few C++ routines of the Hub version of the Gromacs code, in particular, both the header file and the .cpp file of the class "pull" and the source code file "pullutil.cpp" which implement the definitions of the chain reaction coordinate. To use the NP as reference and to define the hydrophobic beads that should form the stalk, the new code makes use of an index file (usually called index.ndx). The latter contains a group named tails, constituted of all the hydrophobic atoms of the membrane (i.e., the lipid tails atoms and the hydrophobic cholesterol atoms) contributing to the reaction coordinate, and a group named NP, constituted by the beads composing the core of the nanoparticle. The chain coordinate parameters are specified via environment variables in a .sh file introduced by Hub et al., that we modified inserting the new parameter *h*.

##### Umbrella sampling simulations

Stalk formation PMFs were computed along the reaction coordinate $\xi_{ch}$ using the umbrella sampling technique. To perform Umbrella Sampling, the starting configuration of each window had to be generated from a pulling simulation in which we slowly pulled the system along $\xi_{ch}$ to drive stalk formation. To do this, we used a harmonic potential (force constant 1000 kJmol−1).

The minimum of the harmonic potential was moved with constant velocity from a minimum value $\xi_{min}$ = 0.25 to a maximum value $\xi_{max}$ =1 in steps of 0.05. The 16 windows were simulated for 1 μs using a harmonic bias potential of 3000 kJ $mol^{-1}$. Then we added sampling near the barrier by re-simulating the window closer to the barrier and introducing two windows at values of $\xi_{ch}$ distant 0.025 from the previously one, using this time a harmonic bias potential of 9000 kJ $mol^{-1}$.

Then, the PMF was constructed from the umbrella histograms with the weighted histogram analysis method (WHAM), as implemented in the g_wham software. This method allows for an estimation of the error on the free energy profile through bootstrap analysis[12].

Furthermore, we also calculated the free energy profiles of stalk destruction, using initial configurations for each window taken from a reverse pulling simulation, in which the system has been brought from the stalk state to the pre-stalk state (from $\xi_{max}$ to $\xi_{min}$).

Figure S2 reports the profiles of stalk formation and destruction for the system with the 4 nm NP. The profiles have been shifted so that the free energy of the pre-stalk state is equal to zero. The profiles of stalk formation and destruction overlap (within error bars), demonstrating the absence of hysteresis in our calculations.

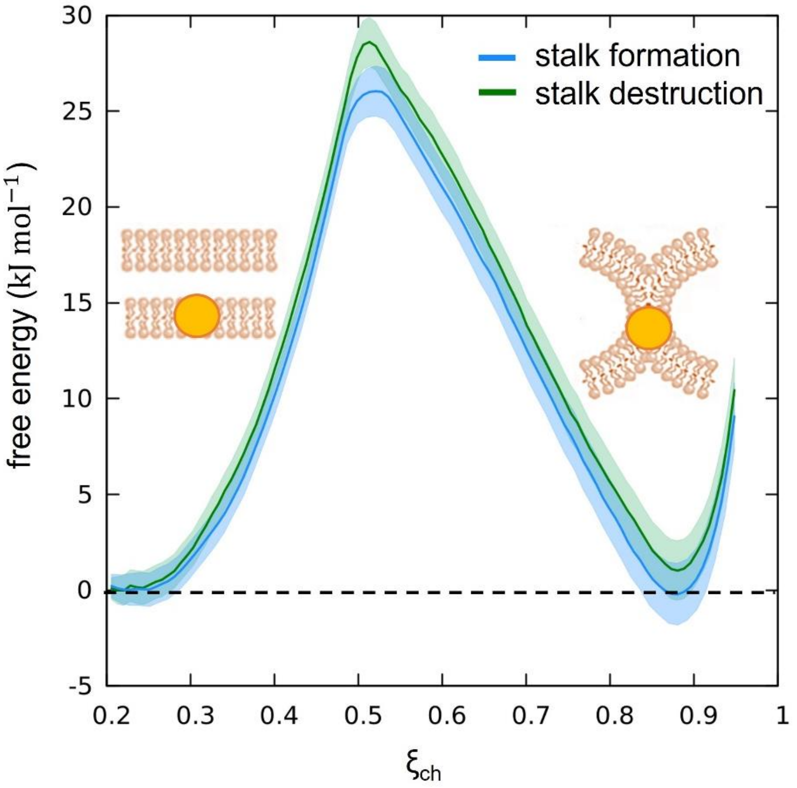


**Figure S2: free energy profiles of stalk formation.** Free energy profiles of stalk formation (light blue) and destruction (green) as a function of $\xi_{ch}$ for NP with a core size of 4 nm. The absence of hysteresis demonstrates the reliability of our CV.

**H-LASA calculation**

For calculating the H-LASA of the NP we created a group composed by all the hydrophobic beads of the ligands (H-MUS) and used it exploiting the *gmx SASA* module of Gromacs. In particular, instead of using a probe of 0.26 nm, that is the radius of a regular Martini bead, we used 0.47 nm, namely the Lennard-Jones sigma of regular Martini interaction. For the case of the embedded NP we considered only the ligands facing the membrane above (see Figure S3 b.). To better compare the different systems, in each case the obtained hydrophobic LASA value was then normalized dividing it by the total LASA (calculated including the MUS charged terminals, MUS-) to obtain the fraction of hydrophobic LASA. Figure S4 shows the time evolution of the H-LASA during a simulation, for NPs of both sizes when embedded in the membrane, while Figure 5 shows the maximum H-LASA, over the simulation time, for the NPs in water.

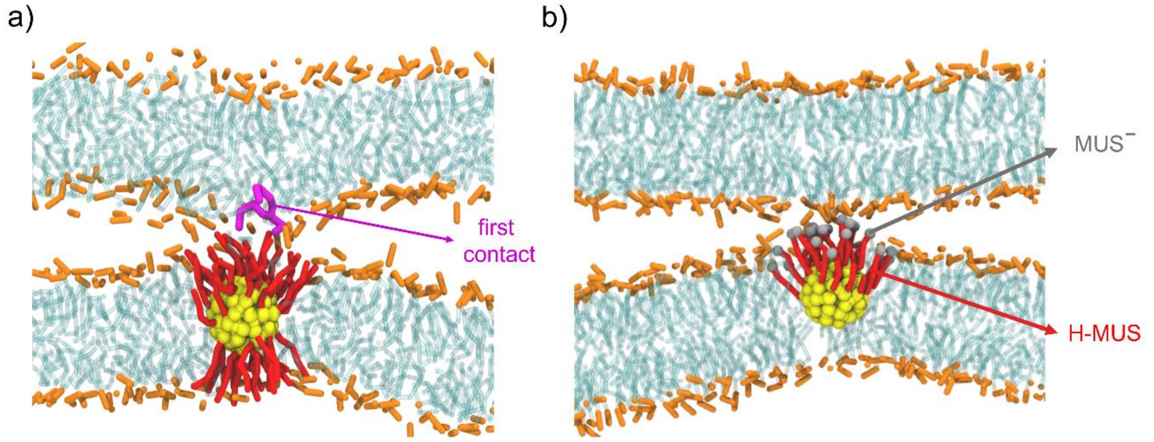


**Figure S3:** a.) Snapshot taken from the system containing a 2nm NP, showing the formation of a hydrophobic contact between a lipid tail of the top bilayer (purple) and the MUS ligands (red). b.) Snapshot in which the beads used to calculate H-LASA are highlighted (in red). They include the OT beads and the hydrophobic beads of the MUS ligands in the upper part of the NP. The charged terminals of the MUS ligands (renamed $MUS^-$) are shown in grey.

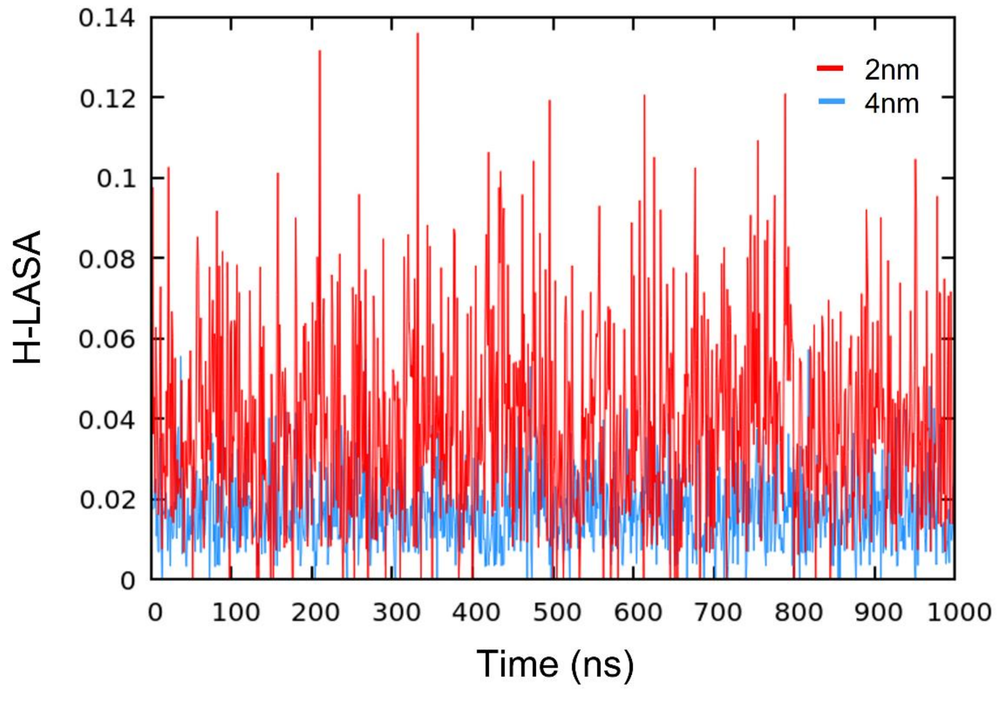


**Figure S4:** time evolution of the H-LASA during a simulation, for NPs of both sizes when embedded in the membrane.

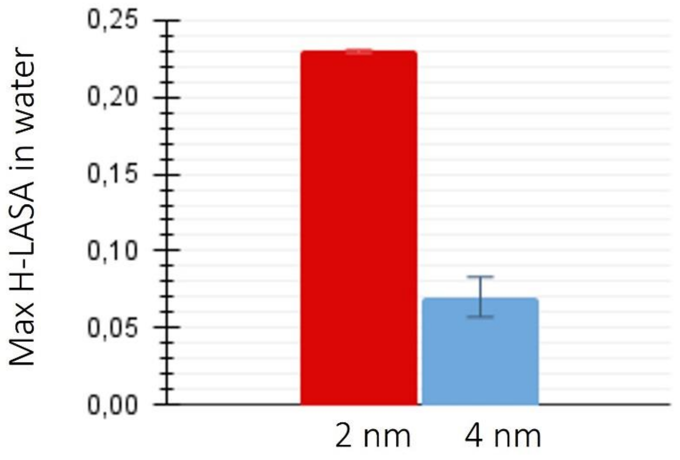


**Figure S5:** Maximum of H-LASA for the two different size NPs in water.

#### Stalk diameter calculation

The total number of hydrophobic atoms of the lipid tails and of cholesterol in the stalk rim was obtained by adding up the atoms located within the 18 slices of the cylinder closer to the NP, and this was made for 10 simulation frames of the umbrella windows with the lower value of $\xi_{ch}$ (the pre-stalk state). The quantities obtained from each frame were then mediated, so average values with the respective standard errors were obtained. For the 2 nm system we obtained the value ${n^{(ap)}}_{2\,nm} = 12.3\ \pm 1.1$ while for the 4 nm one it was obtained ${n^{(ap)}}_{4\,nm} = 6.3\ \pm 0.2$. Since the number of slice $N_s$=23 and the thickness of each slice $ds$=0.1 nm was the same in both cases, also the cylinder height was the same ($N_s \cdot d_s$). We could thus obtain the ratio between the two stalk diameters simply as $\sqrt{\frac{{n^{(ap)}}_{4nm}}{{n^{(ap)}}_{2nm}}}$ founding a value of about 0.7.
